# Beyond Temperature: Environmental Filtering and Functional Shifts in Desert Bird Communities Under Climate Change

**DOI:** 10.1101/2025.10.05.680507

**Authors:** Manasi Mukherjee, Mitali Mukerji, Saran Aadhar

**Author notes:** Corresponding author *Email address:* (Manasi Mukherjee).

## Abstract

Deserts present unique opportunities for understanding biodiversity under climate change, yet global-scale assessments remain rare. Birds serve as sensitive ecological indicators of ecosystem health and biodiversity dynamics. In this study, leveraging crowdsourced bird occurrence records (GBIF) and multi-decadal hydro-meteorological datasets (ERA5, GLEAM), and vegetation data (AVHRR), we show how environmental filtering and functional shifts in avian assemblages drive divergent spatial and temporal species richness patterns across ten major deserts. Noteworthy, responses to warming are distinct, wherein temperature alone cannot universally predict community dynamics. Climatic determinants such as precipitation, soil moisture, and NDVI shaped by greening and browning trends, play region-specific roles. Analyses reveal pronounced guild turnover, with notable declines in top avian predators and complex shifts in omnivores and aquatic predators, exposing ecosystem-level vulnerabilities and altered resilience. These findings underscore the need for a paradigm shift from one-dimensional models to integrative, trait-based approaches to conserve desert biodiversity and function under climate change.

## 1. Introduction

Deserts, comprising nearly one-third of the Earth’s terrestrial surface, are archetypes of ecological resilience, where life persists against the backdrop of extreme temperature fluctuations, persistent aridity, intense solar radiation, and scarce precipitation (Ward, 2010; Maestre et al., 2015). These environments serve as natural laboratories for understanding the biological limits imposed by climate extremes and for examining the mechanisms that allow organisms to withstand and adapt to rapidly changing conditions (Whitford and Duval, 2020; Chown and Gaston, 2008). As the frequency and severity of climatic extremes intensify globally, arid ecosystems are also increasingly recognized for both their vulnerability and their value as sentinel systems for climate change assessment (IPCC, 2023).

Despite their scientific importance, deserts remain among the least studied biomes. Limitations in logistic accessibility, vast and remote landscapes, and the patchy, often elusive nature of desert life forms present considerable challenges to sustained ecological monitoring (Sullivan et al., 2009). Among the biota, birds offer a particularly powerful tool for desert ecological research. Owing to their mobility, visibility, rapid responses to environmental shifts, and well-documented migratory strategies, birds are widely regarded as reliable bioindicators of ecosystem change in arid and as well as non-arid systems (Gregory et al., 2003). Advances in data sharing, citizen science, and trait databases such as Global Biodiversity Information Facility (GBIF) (GBIF.org, 2024) and comprehensive global avian ecological data of AVONET (Tobias et al., 2022)) have enabled researchers to overcome traditional barriers by providing extensive, spatially and temporally explicit data on avian communities (Boakes et al., 2010; Tobias et al., 2022).

( Our previous studies on patterns of bird species richness, migratory categories, and trophic niches across the deserts of the world have revealed pronounced hemispheric and biogeographic contrasts (Mukherjee et al., 2023, 2025; Mukherjee and Mukerji, 2025). The temporal dimension, how these communities and their ecological roles respond to shifting environmental parameters over time, still remain largely underexplored. In particular, it is unclear whether desert bird assemblages mirror the direct impacts of rising temperatures and changing precipitation, or whether their responses are modulated by complex interactions among abiotic drivers (temperature, rainfall, evaporation, soil moisture) and biotic factors (vegetation productivity as indexed by NDVI, resource pulsing, and functional traits) Dai et al., 2022; Han et al., 2024; Wisz et al., 2013). Understanding these patterns is increasingly urgent as deserts experience rapid climate shifts and mounting anthropogenic pressure (Huang et al., 2016; IPCC, 2023). In this study, we assemble and integrate robust crowdsourced and publicly available datasets across ten major desert systems spanning both hemispheres to (i) disentangle the unique temporal climate trajectories of each desert, (ii) assess whether climate variables alone can explain richness and ecological function, and (iii) clarify the deeper links between shifting environmental conditions and dynamic community structures, emphasizing trophic niches and migration. By synthesizing multi-source data and applying contemporary statistical methods, our approach establishes a critical baseline for biodiversity forecasting in Earth’s most extreme and fragile biomes. Our study not only provides foundational knowledge for the conservation of desert biodiversity, but also advances ecological theory by demonstrating the necessity of integrated, trait-based approaches to unravel the impacts of climate variability on ecosystem functionality (Tylianakis et al., 2008; Scheffer et al., 2012).

## 2. Results

### 2.1. Temporal Trends in Avian Species Richness Across Deserts

Analysis of temporal trends in avian species richness across all studied deserts from 2000 to 2020 reveals a consistent, significant increase in species richness for each migratory category (Table 1). The positive slopes of the linear regression models for all categories and deserts confirm that richness has increased steadily and significantly (p < 0.001) over time. Deserts situated near the Tropic of Cancer (Tcan) such as the Arabian, Thar, and Sahara and those close to the Tropic of Capricorn (Tcap) including Great Australian, Kalahari, and Patagonian display marked differences in temporal variation. Tcan deserts generally show higher slope estimates, especially for migratory and partially migratory categories, suggesting a steeper increase in richness, whereas the Tcap deserts show moderate but still significant upward trends. In addition, Tcap deserts demonstrate a prominent and statistically significant increase in sedentary species richness, suggesting that factors driving avian community change extend beyond migratory behaviors and influence resident species as well (Table 1). The differences in slope magnitude and statistical significance underscore variable rates of change between deserts and migratory categories. These differences could possibly be a reflection of the underlying climatic and ecological heterogeneity among these desert systems.

**Table 1:**
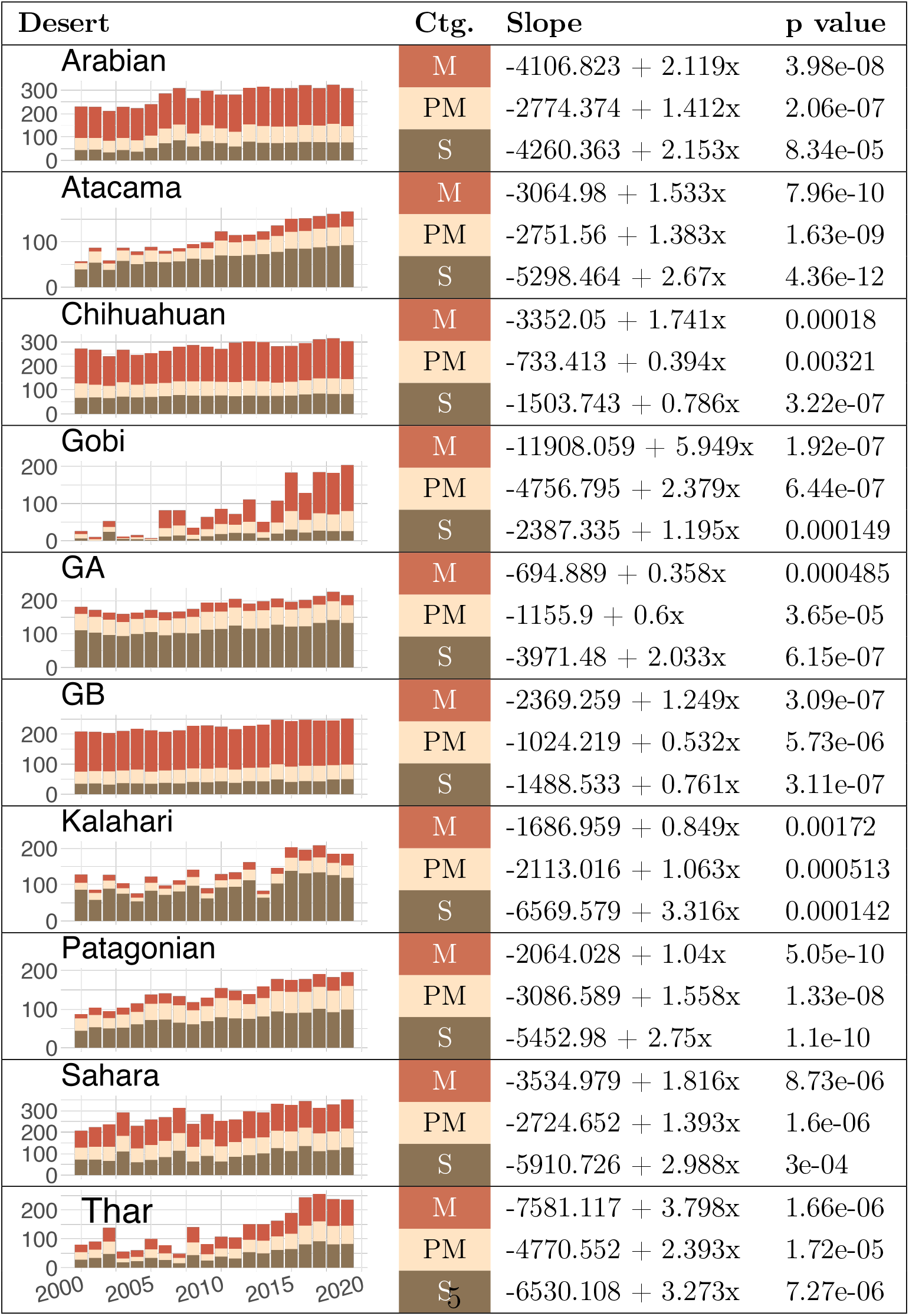
Temporal changes in species richness for avian migratory categories across deserts from 2000-20. The y-axis (Count) indicates annual species richness, while the x-axis shows time in Year. Linear regression slope estimates represent the direction and magnitude of change with time for each migratory category (Ctg.), while p-values indicate statistical significance of trend.

| Desert | Ctg. | Slope | p value |
| --- | --- | --- | --- |
| **Arabian** | M | $-4106.823 + 2.119x$ | 3.98e-08 |
| | PM | $-2774.374 + 1.412x$ | 2.06e-07 |
| | S | $-4260.363 + 2.153x$ | 8.34e-05 |
| **Atacama** | M | $-3064.98 + 1.533x$ | 7.96e-10 |
| | PM | $-2751.56 + 1.383x$ | 1.63e-09 |
| | S | $-5298.464 + 2.67x$ | 4.36e-12 |
| **Chihuahuan** | M | $-3352.05 + 1.741x$ | 0.00018 |
| | PM | $-733.413 + 0.394x$ | 0.00321 |
| | S | $-1503.743 + 0.786x$ | 3.22e-07 |
| **Gobi** | M | $-11908.059 + 5.949x$ | 1.92e-07 |
| | PM | $-4756.795 + 2.379x$ | 6.44e-07 |
| | S | $-2387.335 + 1.195x$ | 0.000149 |
| **GA** | M | $-694.889 + 0.358x$ | 0.000485 |
| | PM | $-1155.9 + 0.6x$ | 3.65e-05 |
| | S | $-3971.48 + 2.033x$ | 6.15e-07 |
| **GB** | M | $-2369.259 + 1.249x$ | 3.09e-07 |
| | PM | $-1024.219 + 0.532x$ | 5.73e-06 |
| | S | $-1488.533 + 0.761x$ | 3.11e-07 |
| **Kalahari** | M | $-1686.959 + 0.849x$ | 0.00172 |
| | PM | $-2113.016 + 1.063x$ | 0.000513 |
| | S | $-6569.579 + 3.316x$ | 0.000142 |
| **Patagonian** | M | $-2064.028 + 1.04x$ | 5.05e-10 |
| | PM | $-3086.589 + 1.558x$ | 1.33e-08 |
| | S | $-5452.98 + 2.75x$ | 1.1e-10 |
| **Sahara** | M | $-3534.979 + 1.816x$ | 8.73e-06 |
| | PM | $-2724.652 + 1.393x$ | 1.6e-06 |
| | S | $-5910.726 + 2.988x$ | 3e-04 |
| **Thar** | M | $-7581.117 + 3.798x$ | 1.66e-06 |
| | PM | $-4770.552 + 2.393x$ | 1.72e-05 |
| | S | $-6530.108 + 3.273x$ | 7.27e-06 |

### 2.2. Temporal Trends in Hydro-meteorological Parameters Across Deserts

The desert and migratory-category wise variable patterns in avian species richness motivated our subsequent exploration of the variability in environmental parameters and ecological functioning across the studied deserts. To examine how temporal variation in the species richness of avian communities in these studied deserts are influenced by their environmental parameters, effect of temperature, precipitation, evaporation, soil moisture and NDVI were analyzed on species richness of each migratory category.

#### 2.2.1. Temperature

Across all deserts studied, mean annual temperature showed a significant rising trend over the years. This increase was most pronounced in the Great Basin, Atacama, Arabian, Patagonian, and Sahara where regression slopes were especially steep, indicating a rapid warming trend. In contrast, the Thar, Great Australian and Kalahari deserts exhibited a relatively slower rate of increase. The positive temperature trend in each desert was statistically significant, as indicated by high R^2^ values and very low p-values, confirming that recent decades have seen substantial and consistent warming throughout these desert systems (Fig. 1A).

**Figure 1:**
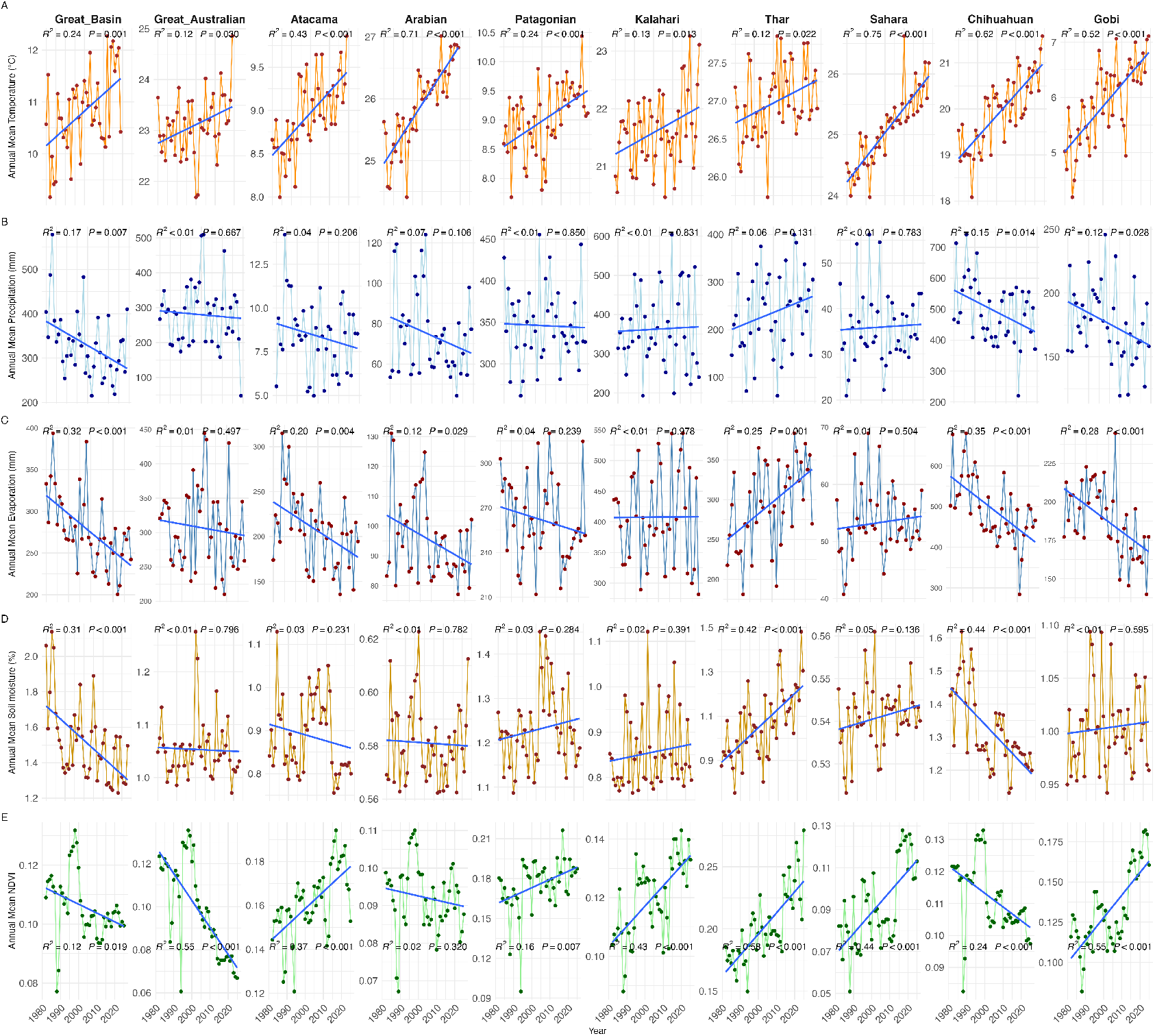
Temporal trends of key hydro-meteorological variables: annual mean temperature (A), precipitation (B), evaporation (C), soil moisture (D), and NDVI (E), across major deserts worldwide. Each sub-panel corresponds to a desert, displaying time series from 1980 to 2020. The linear regression lines (solid smoothers) indicate long-term trends. This plot highlights spatial heterogeneity and temporal variability in climate and vegetation indices critical for understanding desert ecosystem dynamics.

#### 2.2.2. Precipitation

Patterns in annual precipitation remained mostly stable among the deserts with no significant trends. However, deserts like Great Basin, Chihuahuan, Arabian and Gobi showed a significant declining pattern of precipitation (Fig. 1B). These diverse trajectories highlight the heterogeneity in hydrological responses, and generally indicate that changes in precipitation are more desert-specific and less uniform compared to temperature trends.

#### 2.2.3. Evaporation

Annual evaporation generally exhibit variable trends in most deserts, with a few where the slope was negligible. Deserts like Great Basin, Atacama, Arabian, Chihuahuan and Gobi show a decreasing trend of evaporation. The Thar is the only desert that demonstrated especially steep increase in evaporation. (Fig. 1C). These observed increases in evaporation act as amplifiers of local aridity and could potentially accentuate water deficits in sensitive systems.

#### 2.2.4. Soil Moisture

Soil moisture trends diverged sharply across different deserts. Notably, the Great Basin and Chihuahan deserts experienced significant downward slopes, indicating a decline in moisture availability likely driven by increased temperature and evaporation. In contrast, the Arabian, Great Australian, and Gobi deserts appear relatively stable in terms of soil moisture (Fig. 1D). Thar, of all deserts show highest and significant increase in soil moisture trend. These results suggest that for several key regions, declining soil water content is an emerging issue that may have direct ecological impacts, particularly in supporting plant and avian community persistence.

#### 2.2.5. NDVI

NDVI, a proxy for vegetation greenness and productivity, presented both increases and decreases depending on desert location. Deserts such as Sahara, Thar, Kalahari, Gobi and Patagonian deserts exhibit significant temporal increase in NDVI, and a negative trend in the Great Basin, Chihuahuan, Great Australian and Arabian regions (Fig. 1E), highlighting persistent or worsening arid conditions.

Overall, the Great Australian Desert exhibits no significant temporal trends in climatic parameters; however, it shows a significant decline in NDVI over time. In contrast, the Thar Desert, despite a significant increase in evaporation and no corresponding increase in precipitation, maintains high soil moisture levels and demonstrates a significant increasing trend in NDVI. These observations indicate that the effects of rising temperature on desert ecosystems, particularly on vegetation productivity as measured by NDVI, are variable and likely mediated by complex interactions among climatic factors, as illustrated in Figure 1.

### 2.3. Environmental effects on species richness

Based on the climatic variability observed within desert systems, we examined the correlation between major environmental parameters and avian species richness across migratory categories (Fig. 2). The results highlight temperature and NDVI as the most influential determinants of avian species richness in the studied deserts. Species richness of all migratory categories generally showed strong positive correlations with temperature, with notable exceptions in the Gobi, Great Basin, and Thar deserts where the relationships were weak or non-significant. Precipitation and evaporation displayed variable positive and negative correlations, but were not statistically significant predictors of species richness in most deserts, except for migratory categories in the Sahara and Great Australian deserts where significant relationships were observed.

**Figure 2:**
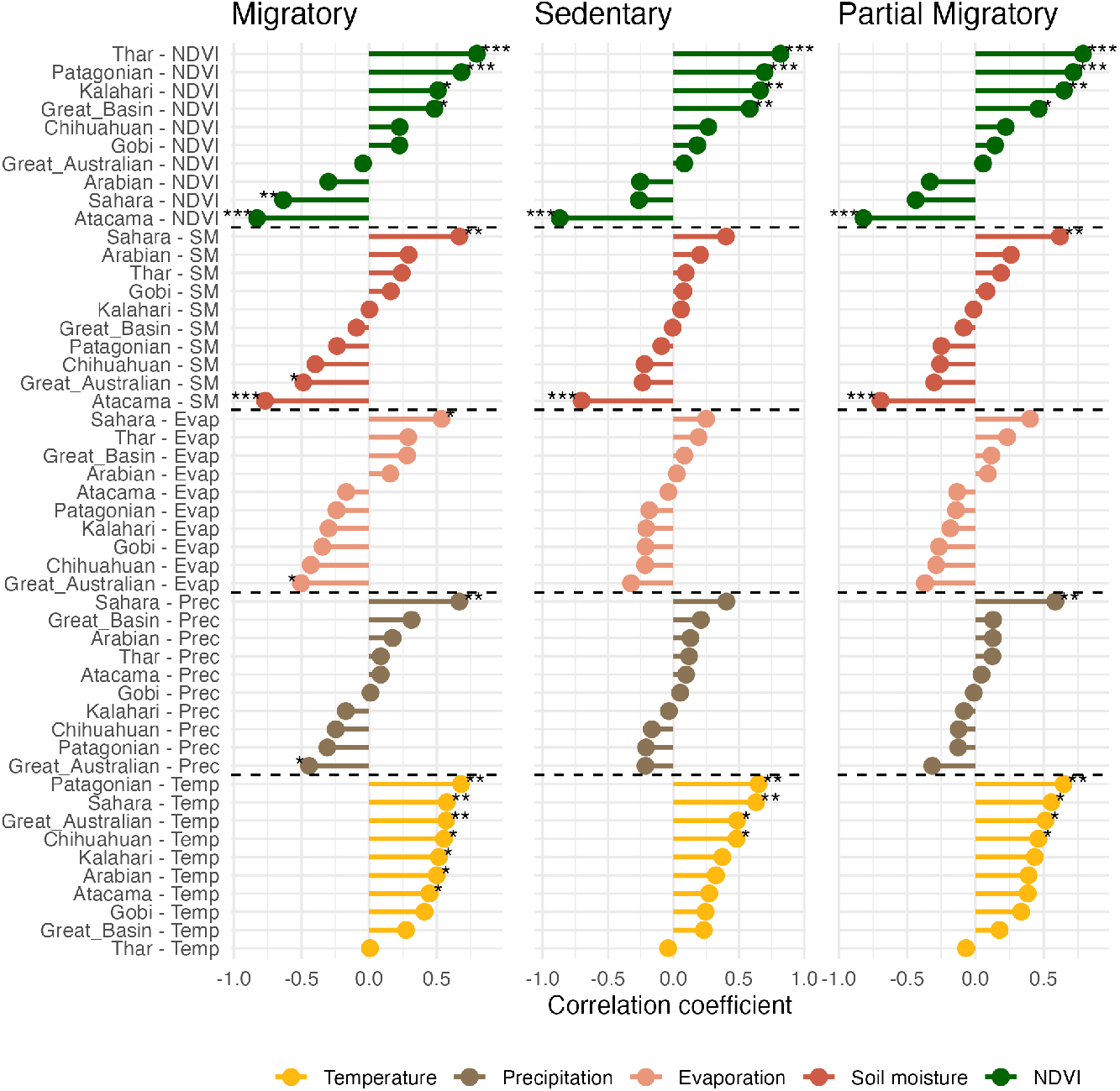
Correlation coefficients between environmental parameters and bird species richness for each of the world’s major deserts, separated by migratory status categories (Migratory, Sedentary, and Partial Migratory species). Each plot displays lollipop lines for five environmental variables, Evaporation, NDVI, Precipitation, Soil Moisture, and Temperature—across individual deserts, with the length and direction of each lollipop indicating the strength and sign of the correlation. Positive values (right of zero) denote a positive association, while negative values indicate a negative association with species richness. Significance is indicated by asterisks:(*** for P < 0.001, ** for P < 0.01 and * for P < 0.05). This plot enables direct comparison of how environmental factors relate to bird species diversity in different deserts and for different movement strategies, highlighting both general patterns and unique local effects.

Distinct patterns emerged for soil moisture over a period of 20 years: Atacama (period average = 0.07 %) exhibited highly significant negative correlations with species richness across all bird categories, whereas the Sahara (period average = 0.04%) displayed highly significant positive correlations, indicating soil moisture as a key driver in these regions. NDVI demonstrated the highest proportion of significant associations, with approximately 70% of deserts showing strong positive correlations with species richness. Positive NDVI correlations were especially notable in the Thar, Patagonian, Kalahari, and Great Basin deserts, while significant negative correlations were found in the Sahara and Atacama for migratory bird categories (Fig. 2). These findings reinforce the dominant role of temperature and NDVI, while emphasizing desert-specific responses to other climatic factors.

While correlation analysis established the relationships between environmental parameters and avian species richness among migratory categories, such relationships may not always be unidirectional or strictly linear, as suggested by temporal variation in the parameters themselves. Thus, the GLM analysis provided more detailed insights into the complex climatic interactions governing species richness patterns in deserts. Results from the 24-year dataset revealed that environmental controls on avian richness are heterogeneous, varying by both desert and migratory category (Table 2). Notably, combinations of environmental predictors explained significant variation in richness for sedentary and partial migratory birds across most deserts, with the exception of the Great Australian Desert, which lacked any significant predictor associations.

**Table 2:**
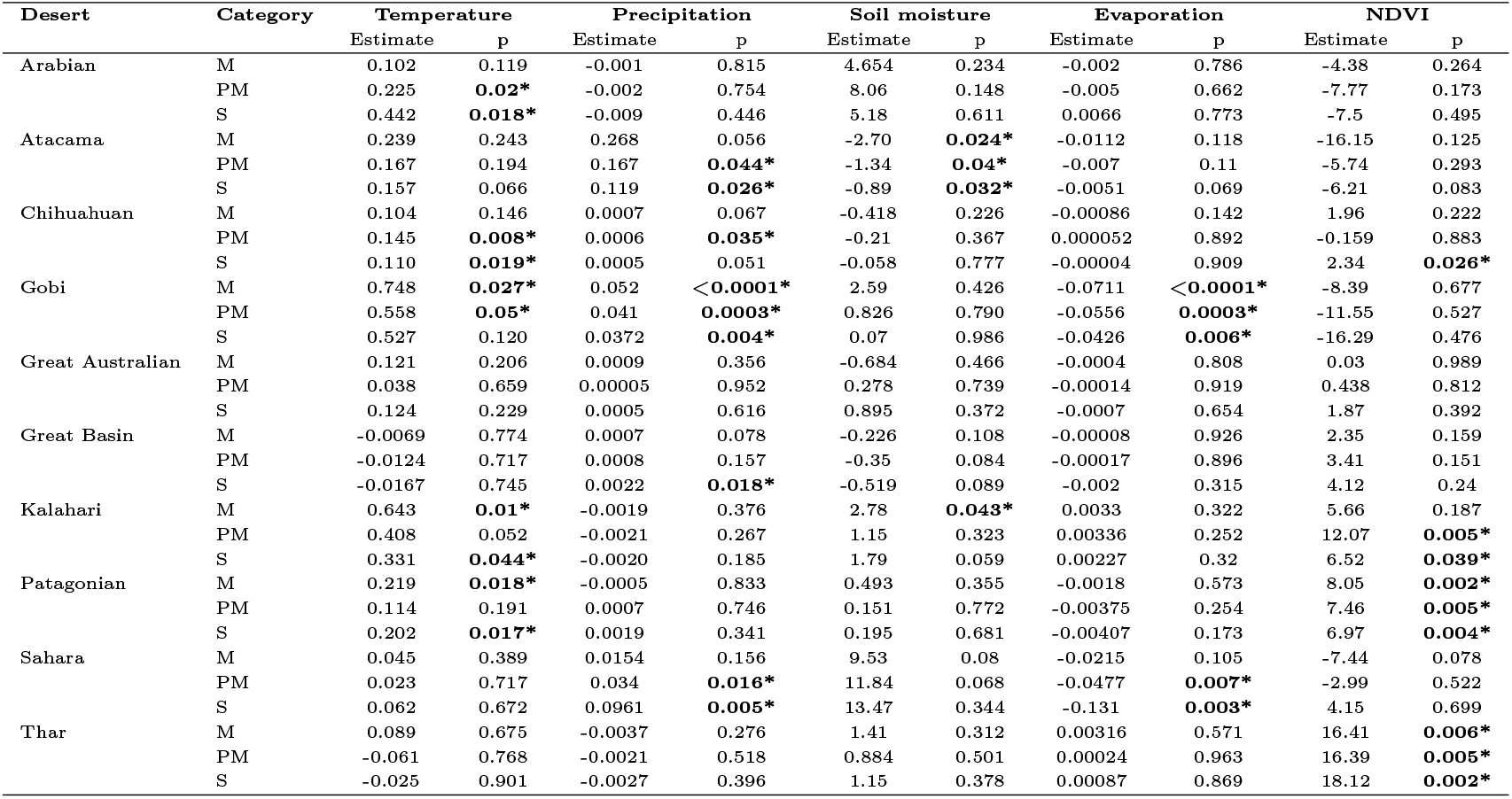
Estimates and p-values from GLM analysis assessing the effects of temperature, precipitation, soil moisture, evaporation, and NDVI on bird species richness across deserts and migratory categories (M = migratory, PM = partial migratory, S = sedentary). Significant p-values (*α* 0.05) are marked with *.

Migratory species richness was generally less explained by the climatic variables tested, indicating that non-climatic factors may exert a stronger influence on this guild. Notably, although correlation analysis indicated that migratory guilds in 70% of the studied deserts were positively associated with temperature, the GLM results identified temperature as a statistically significant determinant only in the Kalahari, Gobi, and Patagonian deserts.

Furthermore, as shown in Figure 2, soil moisture acted as a strong negative predictor of migratory richness in the Atacama Desert, while the GLM also revealed it to be a positive determinant in the Kalahari for this guild (Table 2). Precipitation, which was largely insignificant in the correlation analysis (Figure 2), emerged as a highly significant predictor for sedentary and partial migratory guilds in the Atacama, Gobi, Great Basin, and Sahara deserts in Table 2. In contrast, both the Great Australian and Sahara deserts exhibited weak explanatory power for all environmental parameters, pointing to the potential importance of other, untested ecological or regional variables in these areas. These findings emphasize that the set of most influential environmental factors differs substantially by both desert and migratory category.

### 2.4. Spatio-Temporal variation in Trophic niche of migratory categories

The correlation analysis and GLM results, though did not fully account for the variability in species richness across migratory categories in all deserts, pointed to two major considerations: (i) not all deserts are regulated by the same environmental parameters with regard to guild-wise avian species richness, and (ii) there exists pronounced inter-desert variability in key regulatory factors. Therefore, to better assess the stability and sustainability of these fragile ecosystems under climatic change (Fig. 1), we investigated the temporal patterns in the trophic structure of migratory guilds among recorded avian species.

Over the 23-year assessment period, species richness was classified into ten trophic niches, visualized via a bubble plot (Fig. 3). Across all deserts and migratory categories, invertivores and omnivores were consistently the most abundant trophic niches, a reflection of the arid ecosystems. In particular, sedentary birds exhibited a higher proportion of omnivores, while migratory birds showed a greater relative prominence of invertivores. Significant aquatic predator presence was also notable among migratory birds, especially in the Sahara, Arabian, Great Basin, and Thar.

**Figure 3:**
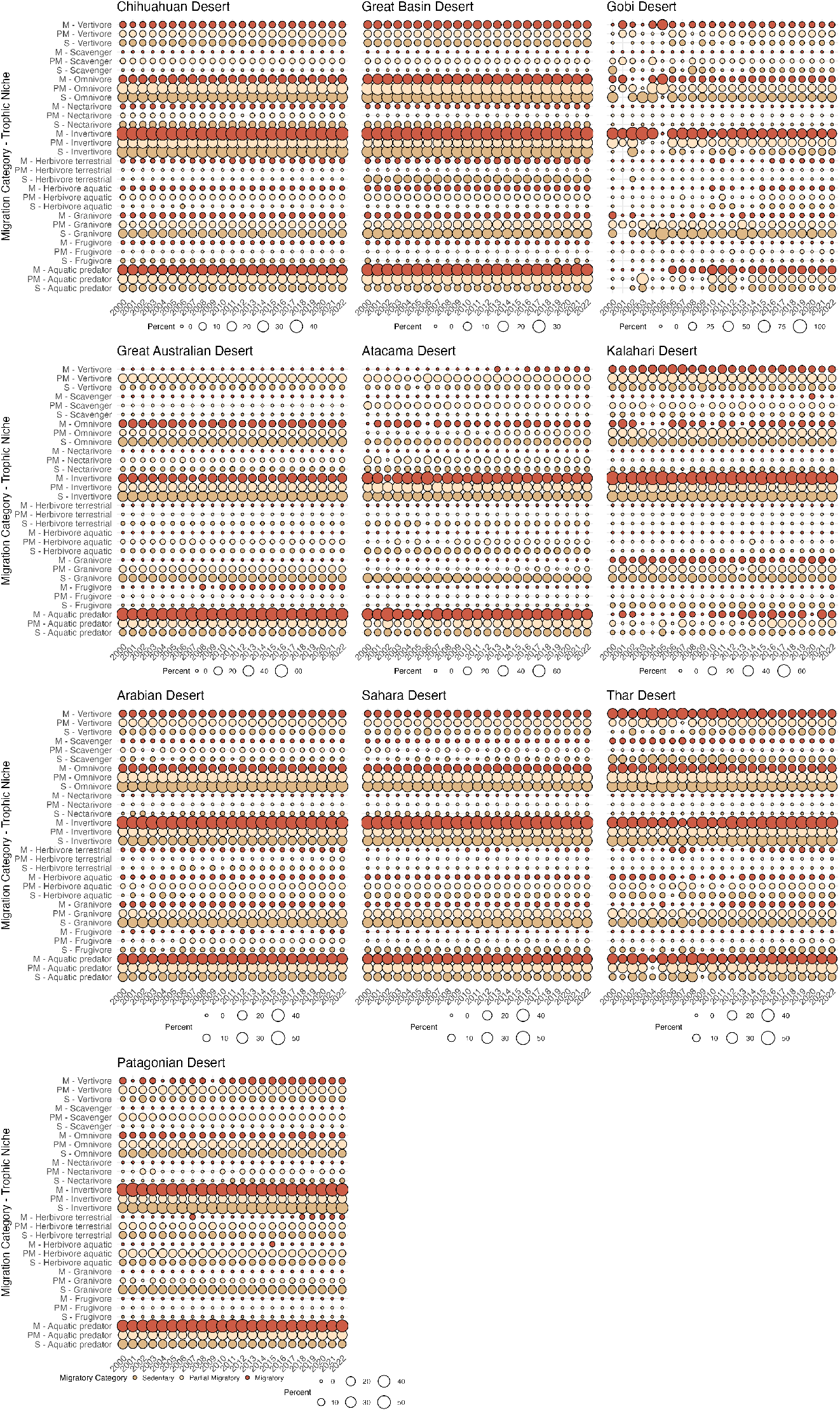
Bubble plots to visualize multidimensional trophic-migratory relationships of Deserts. Trophic niches are mapped along the vertical axis, stratified by migration category, and temporal variation represented along the horizontal axis. Bubble size reflects the proportion of species richness within each niche, while bubble color corresponded to migratory status.

Considerable differences in trophic niche composition were apparent, both among migratory guilds and between deserts. Deserts such as the Chihuahuan, Great Basin, Arabian, Sahara, and Great Australian displayed broad and relatively stable trophic structures across all guilds, indicating stable ecological assemblages with minimal temporal fluctuation. In contrast, Thar, Gobi, and Kalahari deserts demonstrated pronounced temporal variability in trophic guild representation. Kalahari, for instance, exhibited a clear increasing trend in aquatic predator richness among migratory and partial migratory birds. The Gobi desert revealed highly inconsistent trends across nearly all trophic niches, suggesting dynamic shifts in community composition. In the Thar desert, temporal variation in scavenger richness stood out, with pronounced declines among migratory and sedentary scavengers alongside increases in migratory granivores and sedentary frugivores, particularly in the past decade. An analogous divergent pattern was observed for vertivores, wherein migratory vertivores declined, while sedentary vertivores increased in the Thar (Fig. 3).

These findings illustrate that ecological stability and the resilience of avian community trophic structures under changing climates are highly desertand guild-specific, with several deserts exhibiting substantial shifts in resource use and guild dominance over time.

## 3. Discussion

Desert ecosystems represent some of the planet’s most climatically extreme and ecologically complex biomes, exhibiting marked heterogeneity in both environmental drivers and biological responses. Our extended analysis on the spatial patterns previously reported by Mukherjee and Mukerji (2025) across global deserts reveals contrasting temporal patterns in bird species richness segmented by migratory guilds, underscoring fundamental ecological differences between Tcan and Tcap deserts. The relatively balanced species richness among sedentary, partial migratory, and migratory birds in Tcan deserts such as Arabian, Thar, and Sahara likely reflects flexible life histories supported by diverse and heterogeneous habitats adapted to fluctuating environmental conditions (Ma et al., 2023). In contrast, Tcap deserts including Atacama, Patagonian, and Kalahari show notable increases in sedentary species richness, suggesting stronger environmental filtering mechanisms favoring resident birds capable of enduring year-round conditions (Fig. 1).

Climate trends further compound these biogeographic differences. Although rising mean annual temperatures are a near universal feature of the studied deserts (Mamtimin et al., 2011), the rate and nature of warming vary substantially (Fig. 1A). Stark warming trends characterize Arabian and Sahara deserts, while Patagonian and Great Basin deserts experience pronounced internal variability likely tied to latitude and topographical complexity. Deviations evident in the Thar Desert, with comparatively less rising temperatures against increases in precipitation, NDVI, soil moisture, and evaporation (Mishra et al., 2025), emphasize the profound influence of local climatic and ecogeographic context in modulating biome-level responses.

Environmental determinants of avian richness demonstrate complex, context-specific roles. Temperature and NDVI emerge as key influencers, positively correlating with richness across most sites except in harsh systems like Gobi, Great Basin, and Thar, where physiological or ecological constraints potentially limit temperature driven richness enhancement (Fig. 2; Table 2).

These findings echo recent observations that the relationship between heat and avian abundance is not linear globally, especially within latitudinal bands such as 21°–43°, where data have been limited (Kotz et al., 2025). Precipitation and evaporation exert variable, often indirect influences, modulated by regional aridity and hydrological dynamics (Dai et al., 2022; Madouh, 2022); for example, episodic rainfall in hyper-arid Sahara and Atacama does not always translate into ecological benefit (Fig. 2, Table 2) due to rapid desiccation and nutrient-poor soils (Han et al., 2024; Luo et al., 2025; Collins et al., 2025).

The complex interplay of these climate variables is observable through divergent NDVI trends, a proxy for vegetation productivity. While deserts like Patagonian and Thar exhibit “greening” trends aligned with moistening and reduced evaporation, deserts such as Great Australian and Arabian demonstrate “browning” patterns indicative of increasing aridity and vegetation stress. The unique case of Thar Desert’s elevated evaporation yet increasing NDVI may reflect anthropogenic influences, including intensified agriculture, complicating interpretations based solely on natural climatic drivers (Manzoor et al., 2025; Kashyap et al., 2025).

Functional shifts in avian assemblages provide critical insights into ecosystem resilience and vulnerability. Dominance of migratory aquatic predators in Tcap deserts versus migratory scavengers in Tcan deserts (Fig. 3), coupled with guild turnover dynamics in Thar and Gobi deserts, suggest climate-driven disruptions to trophic functions vital for nutrient cycling and ecosystem stability (Anderson et al., 2021). While deserts may appear stable when assessed only through climatic variability (Fig. 1) or trends in species richness (Fig. 1), our results (Fig. 3) show that shifts in biotic assemblages and trophic niche structures reveal underlying fragility in ecosystem function. These changes, which become visible through alterations in guild composition and functional roles, indicate that ecosystem processes can be disrupted despite apparent climatic stability.

In summary, our findings advocate for a paradigm recognizing deserts as individualistic, dynamic systems where species richness and temperature alone inadequately reflects ecological health and functional intactness. Variability in climate trajectories, environmental controls, and trophic assemblages warrants desert-specific interpretation when assessing vulnerability and sustainability under escalating climate change scenarios. This approach aligns with emerging evidence cautioning against broad generalizations and emphasizing localized responses critical for conserving desert biodiversity and ecosystem function. Future research should investigate whether deserts are emerging refugia under climate and land-use change and whether stable abiotic trends conceal critical biotic fragilities. Understanding these under-studied, system-specific dynamics is pivotal for forecasting and mitigating biodiversity loss in Earth’s most extreme environments.

## 4. Conclusion

This study reveals that global desert bird assemblages exhibit diverse and system-specific responses to climate change. Distinct trophic restructuring pathways, with consequential functional shifts threaten critical ecosystem processes such as nutrient cycling and trophic regulation, potentially trigger cascading effects throughout desert food webs. Integrated assessments of environmental and biotic dynamics thus provide vital insights for conserving desert biodiversity and ecosystem services amid accelerating climate change.

## 5. Methods

### 5.1. Desert Selection and Avian Biodiversity Data Curation

This study utilizes avian species richness data for ten major desert biomes across five continents: the Sahara and Kalahari (Africa), Arabian, Thar, and Gobi (Asia), Great Basin and Chihuahuan (North America), Atacama and Patagonian (South America), and the Great Australian Desert (Australia) (Fig. 4). The selection, spatial delineation, and curation of these desert systems and associated bird community data follow the protocol developed in our previous work on the global Avian Atlas (Mukherjee and Mukerji, 2025). Briefly, desert boundaries were defined using spatial polygons based on co-ordinates from (Jung et al., 2020), and avian species richness was extracted from the Global Biodiversity Information Facility (GBIF; GBIF.org 2024) via occurrence data corresponding to those polygons. All avian records including resident, breeding, non-breeding, and passage seasonality categories were considered to encompass the total desert bird community. To mitigate biases inherent in citizen-science-based datasets (such as eBird), rigorous data quality controls were applied as described in our prior study (Mukherjee and Mukerji, 2025), including the exclusion of records with country-coordinate mismatches, invalid timestamps, invalid basis of record and unreliable entries flagged by GBIF.

**Figure 4:**
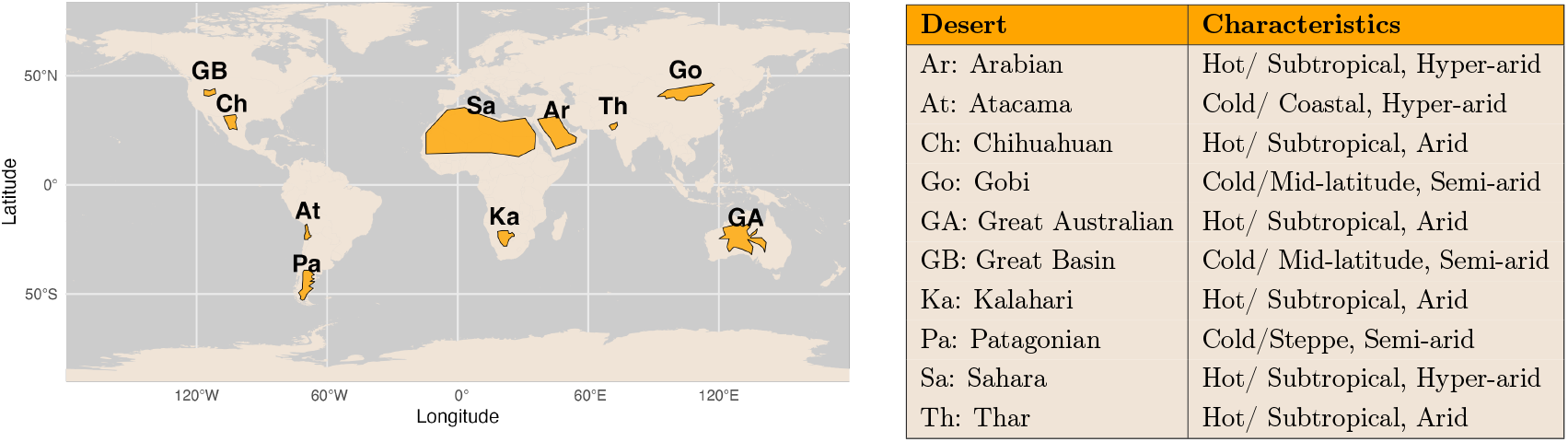
Summary of the characteristics of major deserts under study. Deserts are mapped with orange, and the table beside the map provides the abbreviations used henceforth for the deserts, along with their climatic and ecological characteristics.

### 5.2. Hydro-meteorological and vegetation Data Acquisition and Processing

To examine the relationships between hydro-meteorological variability and avian species richness across the world’s ten largest deserts, we obtained long-term meteorological (Precipitation and Temperature) and hydrological (Evaporation and Soil-moisture) data for each region. Gridded monthly datasets for surface air temperature, total precipitation, evaporation and soil moisture were obtained from the European Centre for Medium-Range Weather Forecasts (ECMWF) reanalysis product (ERA5) (Hersbach et al., 2019), covering the period 1980 to 2022. Normalized Difference Vegetation Index (NDVI), used as a proxy for primary productivity and vegetation cover, was likewise extracted from Advanced Very High Resolution Radiometer (AVHRR) (Stowe et al., 2002), with available records spanning 1980 to 2022.

For each desert, the variables were extracted as area-weighted monthly means, corresponding to the spatial extent of the region as defined by geo-graphic boundaries used in our ecological analyses. To align temporal scales and minimize seasonal bias, all monthly data were aggregated to annual means per desert. In cases of missing or NaN values (resulting from data gaps or conversion artifacts), such entries were excluded before calculating annual statistics, ensuring complete records for temporal trend and regression analyses.

### 5.3. Data Analysis

#### 5.3.1. Temporal Variation in Migratory Category of Birds

We aimed to assess whether observed temporal increases in bird species richness across studied deserts were genuine biological trends or artifacts driven by varying sampling effort over time. To achieve this, we compiled annual species occurrence data for ten deserts from 2000-2020 and summarized the number of unique species (richness) and total observation records (effort) for each year. We then applied generalized linear models (GLMs) with a Poisson error structure to model species richness as a function of year and the number of records, implicit controls for sampling effort. By including effort as a covariate, we accounted for the confounding effect of increased observations potentially inflating richness estimates. The GLM analysis revealed a robust positive association between year and species richness in all deserts, with the coefficient for year being significantly greater than zero, indicating genuine increases in richness over time that cannot be solely attributed to growing sampling effort (Table 3). In contrast, the effect of observational effort (number of records) was generally not significant after accounting for year, suggesting that the observed richness trends were not simply driven by more intensive data collection. These findings provide strong evidence that species richness has increased biologically in these desert ecosystems throughout the study period, independent of sampling bias. The model fit statistics, including Akaike Information Criterion (AIC) and deviance measures, further supported the adequacy of the Poisson GLMs, confirming the validity of our conclusions regarding true temporal changes in bird biodiversity in deserts.

**Table 3:** Generalized linear model (GLM) results for each desert, showing the estimated effects of year and observation effort (number of records) on annual bird species richness from 2000 onward. For each desert, the table reports the model coefficients, standard errors, z-values, and p-values for the year and effort covariates, as well as the Akaike Information Criterion (AIC) as a measure of model fit. Significant year coefficients indicate a genuine increase in species richness over time after accounting for variation in sampling effort.

| Desert | Coefficients |  | Std. error |  | z- value |  | p- value |  | AIC |
| --- | --- | --- | --- | --- | --- | --- | --- | --- | --- |
|  | Year | Effort | Year | Effort | Year | Effort | Year | Effort |  |
| Arabian | 0.026 | -1.43E-06 | 0.004 | 8.63E-07 | 6.056 | -1.655 | 1.4E-09 | 0.098 | 213.2 |
| Atacama | 0.050 | -6.04E-06 | 0.005 | 5.01E-06 | 10.557 | -1.208 | 4.7E-26 | 0.227 | 183.7 |
| Chihuahuan | 0.009 | 3.47E-07 | 0.004 | 6.81E-07 | 2.047 | 0.509 | 4.1E-02 | 0.611 | 194.1 |
| Gobi | 0.118 | -2.20E-05 | 0.005 | 1.02E-05 | 23.058 | -2.168 | 1.2E-117 | 0.030 | 443.9 |
| Great_Aus | 0.014 | 4.94E-07 | 0.004 | 2.10E-06 | 3.578 | 0.235 | 3.5E-04 | 0.814 | 181.6 |
| Great_Basin | 0.011 | 5.18E-08 | 0.004 | 6.56E-07 | 2.449 | 0.079 | 1.4E-02 | 0.937 | 181.3 |
| Kalahari | 0.015 | 6.81E-05 | 0.004 | 8.77E-06 | 4.034 | 7.766 | 5.5E-05 | 0.000 | 271.6 |
| Patagonian | 0.039 | -1.16E-06 | 0.005 | 1.62E-06 | 8.350 | -0.717 | 6.9E-17 | 0.473 | 182.7 |
| Sahara | 0.018 | 1.96E-06 | 0.002 | 1.27E-06 | 7.501 | 1.547 | 6.3E-14 | 0.122 | 227 |
| Thar | 0.047 | 1.46E-05 | 0.005 | 3.52E-06 | 9.682 | 4.156 | 3.6E-22 | 0.000 | 359.8 |

#### 5.3.2. Environmental Effect on Species Richness

To determine the environmental drivers of avian species richness, we analyzed the dataset for each of the ten deserts and three migratory categories (migratory, partial migratory, and sedentary) separately. The relationship between species richness (response variable) and the set of environmental parameters (temperature, precipitation, evaporation, soil moisture, and NDVI) was assessed using Generalized Linear Models (GLMs). A quasipoisson regression with a logarithmic link function was selected for the models to appropriately handle the count-based nature of the species richness data and to account for potential overdispersion. For each model, a p-value of less than 0.05 was considered to indicate a statistically significant relationship. All statistical analyses were conducted in the R programming environment (R Core Team, 2023).

### 5.4. Trophic Niche Analysis and Migratory Behavior Integration

The trophic niche composition of avian communities across the selected desert ecosystems was analyzed to identify patterns in dietary specialization and their association with migratory behaviors. Utilizing the comprehensive dataset by Tobias et al. (2022), species were categorized into ten distinct trophic niches Aquatic Predator, Frugivore, Granivore, Herbivore (Aquatic), Herbivore (Terrestrial), Invertivore, Nectarivore, Omnivore, Scavenger, and Vertivore enabling a detailed examination of ecological roles within each desert group.

To quantify temporal and spatial variation in trophic structure, we calculated the relative abundance (percentage) of species within each trophic niche per desert and year. This approach allowed for the assessment of how dietary preferences fluctuate over time and differ between deserts, providing insight into the dynamic trophic ecology of desert bird assemblages. Furthermore, migratory categories, Sedentary (S), Partial Migratory (PM), and Migratory (M) were integrated into the analysis to probe potential correlations between movement patterns and trophic specialization.

Visualization of these complex interactions was achieved through a series of bubble plots generated in R. In these plots, bubble size reflected the proportion of unique species within each niche, while bubble color corresponded to migratory status, thereby conveying multidimensional trophic-migratory relationships in a single consolidated visualization. Such integrative trophic analyses elucidate the interplay between dietary strategies and migratory behaviors, revealing temporal shifts in niche occupancy and community composition across deserts.

## Authorship contribution statement

**Manasi Mukherjee** contributed in conceptualization, data curation, investigation, methodology, analysis, writing and reviewing. **Mitali Mukerji** contributed in writing and reviewing. **Saran Aadhar** contributed to data curation and editing.

## References

C. B. Anderson, K. Wallen, and J. A. Smith. Unravelling the vertebrate scavenger assemblage in the gobi desert. Journal of Arid Environments, 189:104493, 2021. doi: 10.1016/j.jaridenv.2021.104493.

E. H. Boakes, P. J. McGowan, R. A. Fuller, D. Chang-Qing, N. E. Clark, K. O’Connor, and G. M. Mace. Distorted views of biodiversity: Spatial and temporal bias in species occurrence data. PLoS Biology, 8, 2010. ISSN 15449173. doi: 10.1371/journal.pbio.1000385.

S. L. Chown and K. J. Gaston. Macrophysiology for a changing world, 2008. ISSN 14712970.

S. L. Collins, M. T. Patton, and R. F. Brown. Precipitation-productivity relationships in desert grassland: A test of the double asymmetry hypothesis. Ecology, 106:e70166, 2025. doi: 10.1002/ecy.70166. URL https://esajournals.onlinelibrary.wiley.com/doi/abs/10.1002/ecy.70166.

X. Dai, Z. Yu, A. M. Matheny, W. Zhou, and J. Xia. Increasing evapotranspiration decouples the positive correlation between vegetation cover and warming in the tibetan plateau. Frontiers in Plant Science, 13:974745, 2022. doi: 10.3389/fpls.2022.974745.

GBIF.org. Gbif occurrence download, 3 2024.

R. D. Gregory, D. Noble, R. Field, J. Marchant, M. Raven, and D. W. Gibbons. Using birds as indicators of biodiversity. Methods, 2003.

C. Han, R. Li, and H. Li. Precipitation in july maximizes total above-ground productivity of the desert steppe in inner mongolia, china. PLoS One, 19 (12):e0314983, 2024. doi: 10.1371/journal.pone.0314983.

H. Hersbach, B. Bell, P. Berrisford, G. Biavati, A. Horányi, J. M. Sabater, J. Nicolas, C. Peubey, R. Radu, I. Rozum, D. Schepers, A. Simmons, C. Soci, D. Dee, and J.-N. Thépaut. Era5 monthly averaged data on pressure levels from 1979 to present [dataset]. copernicus climate change service (c3s) climate data store (cds), 2019.

J. Huang, H. Yu, X. Guan, G. Wang, and R. Guo. Accelerated dryland expansion under climate change. Nature Climate Change, 6, 2016. ISSN 17586798. doi: 10.1038/nclimate2837.

IPCC. Summary for policymakers. in: Climate change 2023: Synthesis report. contribution of working groups i, ii and iii to the sixth assessment report of the intergovernmental panel on climate change. Technical report, 2023.

M. Jung, P. R. Dahal, S. H. Butchart, P. F. Donald, X. D. Lamo, M. Lesiv, V. Kapos, C. Rondinini, and P. Visconti. A global map of terrestrial habitat types. Scientific Data, 7, 2020. ISSN 20524463. doi: 10.1038/s41597-020-00599-8.

R. Kashyap, J. Kuttippurath, and V. K. Patel. Agriculture intensification and moisture-induced thar desert greening : implications for energy balance, socio-economy, and biodiversity. GIScience & Remote Sensing, 62:2483458, 2025. doi: 10.1080/15481603.2025.2483458. URL https://doi.org/10.1080/15481603.2025.2483458.

M. Kotz, T. Amano, and J. E. M. Watson. Large reductions in tropical bird abundance attributable to heat extreme intensification. Nature Ecology & Evolution, 2025. ISSN 2397-334X. doi: 10.1038/s41559-025-02811-7. URL https://doi.org/10.1038/s41559-025-02811-7.

Y. Luo, J. Cheng, Z. Cao, H. Zhang, P. Danba, J. Wang, Y. Wang, R. Zhang, C. Zhang, Y. Feng, and S. Wei. Effect of precipitation change on desert steppe aboveground productivity. Biology (Basel), 14(8): 1010, 2025. doi: 10.3390/biology14081010.

L. Ma, S. R. Conradie, C. L. Crawford, A. S. Gardner, M. R. Kearney, I. M. D. Maclean, A. E. McKechnie, C.-R. Mi, R. A. Senior, and D. S. Wilcove. Global patterns of climate change impacts on desert bird communities. Nature Communications, 14(1): 211, 2023. ISSN 2041-1723. doi: 10.1038/s41467-023-35814-8. URL https://doi.org/10.1038/s41467-023-35814-8.

T. A. Madouh. Eco-physiological responses of native desert plant species to drought and nutritional levels: Case of kuwait. Frontiers in Environmental Science, 10, 2022. ISSN 2296665X. doi: 10.3389/fenvs.2022.785517.

F. T. Maestre, M. Delgado-Baquerizo, T. C. Jeffries, D. J. Eldridge, V. Ochoa, B. Gozalo, J. L. Quero, M. García-Gómez, A. Gallardo, W. Ulrich, M. A. Bowker, T. Arredondo, C. Barraza-Zepeda, D. Bran, A. Florentino, J. Gaitán, J. R. Gutiérrez, E. Huber-Sannwald, M. Jankju, R. L. Mau, M. Miriti, K. Naseri, A. Ospina, I. Stavi, D. Wang, N. N. Woods, X. Yuan, E. Zaady, and B. K. Singh. Increasing aridity reduces soil microbial diversity and abundance in global drylands. Proceedings of the National Academy of Sciences of the United States of America, 112, 2015. ISSN 10916490. doi: 10.1073/pnas.1516684112.

B. Mamtimin, A. M. Et-Tantawi, D. Schaefer, F. X. Meixner, and M. Domroes. Recent trends of temperature change under hot and cold desert climates: Comparing the sahara (libya) and central asia (xinjiang, china). Journal of Arid Environments, 75, 2011. ISSN 01401963. doi: 10.1016/j.jaridenv.2011.06.007.

A. T. Manzoor, N. Habib, and S. Abbas. Performance of ndvi and gosif in monitoring vegetation responses to rainfall in a desert ecosystem. Remote Sensing Applications: Society and Environment, 39: 101621, 2025. ISSN 2352-9385. doi: 10.1016/j.rsase.2025.101621. URL https://www.sciencedirect.com/science/article/pii/S2352938525001740.

V. Mishra, H. Solanki, and R. Nemani. Greening of the Thar Desert driven by climate change and human interventions. Cell Reports Sustainability, 2(5): 100364, 2025. ISSN 2949-7906. doi: 10.1016/j.crsus.2025.100364. URL https://www.sciencedirect.com/science/article/pii/S2949790625000606.

M. Mukherjee and M. Mukerji. Avian atlas: Unveiling the diversity divide in major global desert realms. Ecological Indicators, 171:113094, 2025. ISSN 1470-160X. doi: 10.1016/j.ecolind.2025.113094. URL https://www.sciencedirect.com/science/article/pii/S1470160X25000238.

M. Mukherjee, A. Paul, and M. Mukerji. Biotic assessment of crowdsourced data defines four ecoregions in Thar: A novel approach for community engagement in conservation. Global Ecology and Conservation, 46:e02559, 2023. ISSN 2351-9894. doi: 10.1016/j.gecco.2023.e02559.

M. Mukherjee, D. Banerjee, I. Sharma, and M. Mukerji. Wastelands or preferred-lands? indicators for redefining desert conservation. Journal of Arid Environments, 229:105371, 2025. ISSN 0140-1963. doi: 10.1016/j.jaridenv.2025.105371. URL https://www.sciencedirect.com/science/article/pii/S0140196325000552.

R Core Team. R: A language and environment for statistical computing. R Foundation for Statistical Computing, Vienna, Austria, 2023. URL https://www.R-project.org/.

M. Scheffer, S. R. Carpenter, T. M. Lenton, J. Bascompte, W. Brock, V. Dakos, J. V. D. Koppel, I. A. V. D. Leemput, S. A. Levin, E. H. V. Nes, M. Pascual, and J. Vandermeer. Anticipating critical transitions, 2012. ISSN 10959203.

L. L. Stowe, H. Jacobowitz, G. Ohring, K. R. Knapp, and N. R. Nalli. The advanced very high resolution radiometer (avhrr) pathfinder atmosphere (patmos) climate dataset: Initial analyses and evaluations. Journal of Climate, 15, 2002. ISSN 08948755. doi: 10.1175/1520-0442(2002)015<1243:TAVHRR>2.0.CO;2.

B. L. Sullivan, C. L. Wood, M. J. Iliff, R. E. Bonney, D. Fink, and S. Kelling. ebird: A citizen-based bird observation network in the biological sciences. Biological Conservation, 142, 2009. ISSN 00063207. doi: 10.1016/j.biocon.2009.05.006.

J. A. Tobias, C. Sheard, A. L. Pigot, A. J. M. Devenish, J. Yang, F. Sayol, M. H. C. Neate-Clegg, N. Alioravainen, T. L. Weeks, R. A. Barber, P. A. Walkden, H. E. A. MacGregor, S. E. I. Jones, C. Vincent, A. G. Phillips, N. M. Marples, F. A. Montaño-Centellas, V. Leandro-Silva, S. Claramunt, B. Darski, B. G. Freeman, T. P. Bregman, C. R. Cooney, E. C. Hughes, E. J. R. Capp, Z. K. Varley, N. R. Friedman, H. Korntheuer, A. Corrales-Vargas, C. H. Trisos, B. C. Weeks, D. M. Hanz, T. Töpfer, G. A. Bravo, V. Remeš, L. Nowak, L. S. Carneiro, A. J. Moncada R. B. Matysioková, D. T. Baldassarre, A. Martínez-Salinas, J. D. Wolfe, P. M. Chapman, B. G. Daly, M. C. Sorensen, A. Neu, M. A. Ford, R. J. Mayhew, L. Fabio Silveira, D. J. Kelly, N. N. D. Annorbah, H. S. Pollock, A. M. Grabowska-Zhang, J. P. McEntee, J. Carlos T. Gonzalez, C. G. Meneses, M. C. Muñoz, L. L. Powell, G. A. Jamie, T. J. Matthews, O. Johnson, G. R. R. Brito, K. Zyskowski, R. Crates, M. G. Harvey, M. Jurado Zevallos, P. A. Hosner, T. Bradfer-Lawrence, J. M. Maley, F. G. Stiles, H. S. Lima, K. L. Provost, M. Chibesa, M. Mashao, J. T. Howard, E. Mlamba, M. A. H. Chua, B. Li, M. I. Gómez, N. C. García, M. Päckert, J. Fuchs, J. R. Ali, E. P. Derryberry, M. L. Carlson, R. C. Urriza, K. E. Brzeski, D. M. Prawiradilaga, M. J. Rayner, E. T. Miller, R. C. K. Bowie, R.-M. Lafontaine, R. P. Scofield, Y. Lou, L. Somarathna, D. Lepage, M. Illif, E. L. Neuschulz, M. Templin, D. M. Dehling, J. C. Cooper, O. S. G. Pauwels, K. Analuddin, J. Fjeldså, N. Seddon, P. R. Sweet, F. A. J. DeClerck, L. N. Naka, J. D. Brawn, A. Aleixo, K. Böhning-Gaese, C. Rahbek, S. A. Fritz, G. H. Thomas, and M. Schleuning. Avonet: morphological, ecological and geographical data for all birds. Ecology Letters, 25(3): 581–597, 2022. doi: 10.1111/ele.13898. URL https://onlinelibrary.wiley.com/doi/abs/10.1111/ele.13898.

J. M. Tylianakis, R. K. Didham, J. Bascompte, and D. A. Wardle. Global change and species interactions in terrestrial ecosystems, 2008. ISSN 1461023X.

D. Ward. The Biology of Deserts. Oxford University Press, apr 2010. ISBN 9780191728143. doi: 10.1093/ACPROF:OSO/9780199211470.001.0001.

W. G. Whitford and B. D. Duval. Ecology of Desert Systems. 2020. doi: 10.1016/C2017-0-02227-9.

M. S. Wisz, J. Pottier, W. D. Kissling, L. Pellissier, J. Lenoir, C. F. Damgaard, C. F. Dormann, M. C. Forchhammer, J. A. Grytnes, A. Guisan, R. K. Heikkinen, T. T. Høye, I. Kühn, M. Luoto, L. Maiorano, M. C. Nilsson, S. Normand, E. Öckinger, N. M. Schmidt, M. Termansen, A. Timmermann, D. A. Wardle, P. Aastrup, and J. C. Svenning. The role of biotic interactions in shaping distributions and realised assemblages of species: Implications for species distribution modelling. Biological Reviews, 88, 2013. ISSN 14647931. doi: 10.1111/j.1469-185X.2012.00235.x.

